# E6 proteins from high-risk HPV, low-risk HPV, and animal papillomaviruses activate the Wnt/β-catenin pathway through E6AP-dependent degradation of NHERF1

**DOI:** 10.1101/518282

**Authors:** Camille M. Drews, Samuel Case, Scott B. Vande Pol

## Abstract

High-risk human papillomavirus (HPV) E6 proteins associate with the cellular ubiquitin ligase E6-Associated Protein (E6AP), and then recruit both p53 and certain cellular PDZ proteins for ubiquitination and degradation by the proteasome. Low-risk HPV E6 proteins also associate with E6AP, yet fail to recruit p53 or PDZ proteins; their E6AP-dependent targets have so far been uncharacterized. We found a cellular PDZ protein called Na+/H+ Exchanger Regulatory Factor 1 (NHERF1) is targeted for degradation by both high and low-risk HPV E6 proteins as well as E6 proteins from diverse non-primate mammalian species. NHERF1 was degraded by E6 in a manner dependent upon E6AP ubiquitin ligase activity but independent of PDZ interactions. A novel structural domain of E6, independent of the p53 recognition domain, was necessary to associate with and degrade NHERF1, and the NHERF1 EB domain was required for E6-mediated degradation. Degradation of NHERF1 by E6 activated canonical Wnt/β-catenin signaling, a key pathway that regulates cell growth and proliferation. Expression levels of NHERF1 increased with increasing cell confluency. This is the first study in which a cellular protein has been identified that is targeted for degradation by both high and low-risk HPV E6 as well as E6 proteins from diverse animal papillomaviruses. This suggests that NHERF1 plays a role in regulating squamous epithelial growth and further suggests that the interaction of E6 proteins with NHERF1 could be a common therapeutic target for multiple papillomavirus types.

**Author summary:** Papillomaviruses cause benign squamous epithelial tumors through the action of virally encoded oncoproteins termed E6 and E7, which are classified as either high or low-risk based upon the propensity of the tumor to evolve into cancer. E6 proteins from both high and low-risk HPVs interact with a cellular ubiquitin ligase called E6AP. High-risk E6 proteins hijack E6AP ubiquitin ligase activity to target p53 for degradation. Degradation targets of the low-risk E6 proteins in complex with E6AP have not been described. Here, we describe a protein called NHERF1 that is targeted for degradation by both high and low-risk E6 proteins, as well as E6 proteins from diverse animal species. Degradation of NHERF1 resulted in activation of an oncogenic cellular signaling pathway called Wnt. Identification of NHERF1 as a highly conserved E6 degradation target could inform therapies directed against both low-risk HPVs and cancer-inducing high-risk HPVs.

## Introduction

Human papillomaviruses (HPVs) are small DNA tumor viruses that cause squamous epithelial papillomas in which the virus replicates. The papillomas are initially benign and the host is usually able to clear the underlying HPV infection over time. However, a subset of HPV infections may result in lesions that persist and grow to harmful size or that have a propensity to evolve into carcinomas [1]. The cancer-causing HPV types are called high-risk and the most commonly occurring high-risk types are HPV16 and HPV18. Worldwide, high-risk HPVs are responsible for 5% of cancers, with cervical cancer being the most common [2]. HPV types that are not associated with malignancies are termed low-risk HPV; although low-risk for malignancies, the size and location of the benign papillomas can render these lesions medically serious [3].

Beyond HPVs, papillomaviruses have been isolated from mammalian species including rodents, primates, bats, cetaceans, and ungulates [4], and are clustered into related genera based upon the divergence of the L1 capsid protein nucleotide sequence (both high and low-risk mucosal HPV types discussed in this study belong to the primate Alpha genera) [5]. Most non-human papillomaviruses encode E6 proteins that are similar in predicted fold to high-risk HPV16 E6 [6]. When diverse mammalian papillomaviruses are clustered based on their E6 sequence similarity, two main groups of papillomaviruses emerge: those that encode E6 proteins that bind to the Notch co-activator Mastermind Like 1 (MAML1), and those that bind to a cellular E3 ubiquitin ligase called E6-Associated Protein (E6AP) [7]. An E6 protein that preferentially binds MAML1 suppresses MAML1 transcriptional activation, while an E6 protein that preferentially binds E6AP stimulates E6AP E3 ubiquitin ligase activity to then target additional cellular proteins recruited by E6 to E6AP for ubiquitination and degradation by the proteasome [7].

The difference between the propensity of high and low-risk HPVs to cause cancer is secondary to differences between their respective E6 and E7 oncoproteins [8]. E6 and E7 from both high and low-risk HPVs bind cellular E3 ubiquitin ligases and hijack their ubiquitin ligase activity to perturb certain cellular proteins that are recruited by E6 or E7 [9]. Both high-risk and low-risk E7 proteins interact with ubiquitin ligases of the cullin and N-end rule families and target the degradation of additional cellular proteins recruited to E7 such as pocket-family proteins (RB, RBL1, and RBL2) and PTPN14 [10, 11]. High and low-risk E7 proteins target certain cellular proteins in common (such as the RBL2 p130 pocket protein) [12]. However only high-risk HPV E7 types interact with and target the degradation of the retinoblastoma (RB) pocket protein [13, 14], which has implications for the carcinogenic properties of high-risk E7. High and low-risk Alpha genera HPV E6 proteins interact with the cellular E3 ubiquitin ligase E6AP [15–17], but only cellular proteins targeted for degradation by the high-risk E6 protein (such as p53) are well established [18, 19].

Another striking difference between high and low-risk E6 is the presence of a PDZ binding motif (PBM) at the extreme carboxy terminus of high-risk E6 proteins [20–22]}. The high-risk PBM enables E6 to interact with a group of cellular proteins termed PDZ proteins, all of which contain PDZ (PSD-90/Dlg1/ZO-1) homology domains [23]. The targeted degradation of cellular proteins that are recruited through interaction with the high-risk E6 PBM has been controversial, but the E6 PBM functionally promotes retention of the viral DNA plasmid within infected cells [24]; the E6 PBM function can be rescued by disruption of p53 function [25]. Although low-risk E6 proteins bind E6AP, they do not have a PBM at the carboxy-terminus [20], do not interact with p53 [17], and no cellular targets of the low-risk E6+E6AP complex have been described. Such cellular targets would be presumed to be of exceptional interest since they would be common to both high and low-risk E6 proteins, just as RBL2 is a common target of the high and low-risk HPV E7 protein.

In this study, we identify the PDZ-adapter protein NHERF1 as degraded by both high and low-risk E6 proteins, in a manner dependent upon the ubiquitin ligase activity of E6AP and the proteasome. Other E6 proteins from diverse species where E6 could bind E6AP were also able to initiate NHERF1 degradation, indicating the conservation of this function. Interaction of NHERF1 with E6 required prior association of E6 with E6AP, and we identified a novel interaction domain within 16E6 that is required. Finally, the targeted degradation of NHERF1 by both low and high-risk E6 proteins resulted in the activation of canonical Wnt signaling, connecting the degradation of NHERF1 by E6 to the activation of an oncogenic signaling pathway.

## Results

### NHERF1 protein levels are reduced by high and low-risk E6 proteins in the presence of E6AP_WT

NHERF1 was previously shown to be degraded by HPV16 E6 (16E6) (but not by HPV18 E6 (18E6) or HPV11 E6 (11E6)) through an interaction requiring the PBM of 16E6 [26]. In our proteomic studies of cellular proteins that associate with the 16E6 and 18E6 PBMs [27], we did not identify NHERF1, but in other experiments observed a reduction of NHERF1 protein levels by 16E6, 18E6, and 11E6. In order to characterize the reduction of NHERF1 by these E6 proteins, we performed transient transfections into E6AP-null 8B9 cells reconstituted with either WT E6AP (E6AP_WT) or a mutant E6AP defective in ubiquitin ligase activity (E6AP_Ub^−^). E6APs were co-transfected with plasmids encoding p53, NHERF1, and 16E6, 16E6 deleted of the PBM (16E6ΔPBM), 11E6, or 18E6. Consistent with published literature, p53 was degraded by high-risk E6 proteins (16E6 and 18E6) independently of a PBM [25] and dependent upon E6AP ubiquitin ligase activity [28] (Fig 1). Expression of 11E6 together with E6AP_WT resulted in a lack of p53 degradation by low-risk E6 (11E6), corroborating published findings [17, 29]. However, NHERF1 protein levels were reduced by each observed E6 protein (Fig 1A), in contrast to what has been previously published [26].

**Fig 1.**
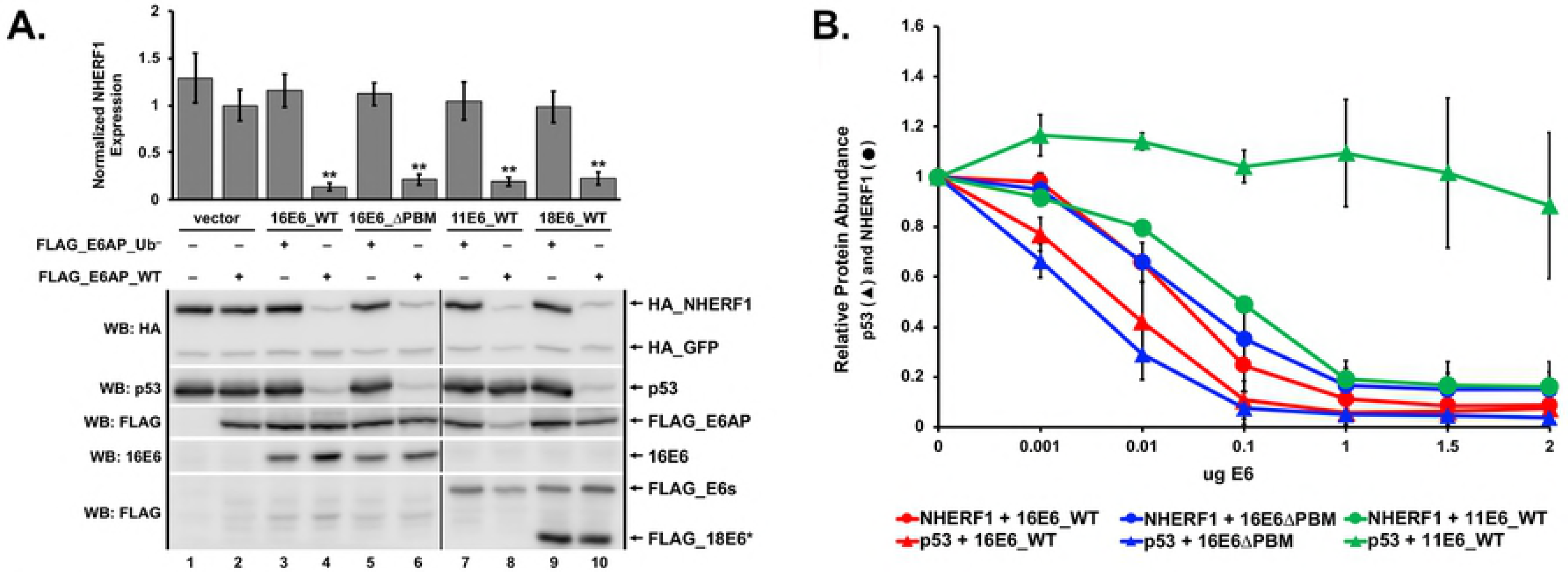
NHERF1 protein levels are reduced by both high and low-risk E6 proteins. (A) NHERF1 protein levels are reduced in an E6 and E6AP dependent manner. Plasmids encoding the indicated FLAG_E6AP (1 ug), HA_NHERF1 (0.5 ug), human p53 (0.5 ug), HA_GFP (0.08 ug), and the listed E6 proteins (1 ug) were transiently transfected into E6AP-null 8B9 cells and HA-NHERF1 expression was analyzed by western blot. Reduction of NHERF1 protein levels by high or low-risk E6 requires ligase active E6AP (E6AP_WT) but does not require the E6 PDZ binding motif (PBM). FLAG_18E6* is a truncated splice isoform of 18E6. E6AP_Ub^−^denotes an E6AP mutant defective for ubiquitin ligase activity. Quantitation is the result of three independent experiments (N=3) where NHERF1 levels are normalized to co-transfected HA_GFP. Shown is a single representative blot. Vertical black line in blots represents removal of an irrelevant sample. The means of triplicate independent experiments ± standard error are shown. N=3. *<0.05, **<0.01 by Student’s t-test. (B) Reduction of NHERF1 protein is not an overexpression artifact. Titrations of the indicated E6 proteins were co-transfected with FLAG_E6AP_WT (1 ug), HA_GFP (0.02 ug), and either HA_NHERF1 (0.5 ug) or p53 (0.5 ug) in murine 8B9 cells. With increased E6 expression, NHERF1 decreased for each E6 protein parallel with p53. As expected, p53 degradation was observed for the high-risk 16E6 proteins (both WT and ΔPBM) but not by low-risk 11E6 protein despite reduction of NHERF1 protein levels by 11E6. The means of triplicate independent experiments ± standard error are shown.

To ensure the reduction of NHERF1 by either high or low-risk E6 proteins was not due to an overexpression artifact, we performed an E6 titration experiment (Fig 1B). Representative western blots from which the quantification in Fig 1B was derived are shown in S1 Fig. Three different E6 proteins were used: 16E6_WT, 16E6ΔPBM, and 11E6_WT. We used p53 as a control for 16E6-mediated degradation, as multiple studies have shown low-risk E6 proteins (11E6) do not degrade p53 [17, 29] while high-risk 16E6 is able to degrade p53 independent of the presence of a PBM [25]. Observing the degradation of p53 in cells expressing variable amounts of E6 provided a guide for physiologically relevant E6 expression levels. NHERF1 and p53 protein levels were similarly reduced by both 16E6_WT and 16E6ΔPBM at the various E6 titrations (Fig 1B). 11E6_WT was unable to initiate the degradation of p53 but targeted NHERF1 at levels similar to those required by 16E6. Deletion of the 16E6 PBM did not impact the degradation of p53 or reduction of NHERF1 protein levels by 16E6.

### NHERF1 protein levels are sensitive to cell confluency

We established that NHERF1 protein levels are reduced by E6 in a transient transfection system. To determine whether low levels of stable 16E6-expression could initiate the reduction of NHERF1 protein levels, we retrovirally transduced normal immortalized keratinocytes with either empty vector or 16E6_WT and observed NHERF1 protein levels. Initially, our results were variable. We hypothesized that keratinocyte confluency may affect NHERF1 protein levels. To test this possibility, we seeded vector-transduced and 16E6_WT-transduced keratinocytes at three different cell densities: 5×10^3^ cells/cm^2^ (very sub-confluent), 1.3×10^4^ cells/cm^2^ (sub-confluent), and 2.6×10^4^ cells/cm^2^ (mid-confluent). NHERF1 protein levels increased with an increase in cell density and 16E6_WT consistently reduced NHERF1 (Fig 2A). In order to determine if changes in NHERF1 levels with confluency were secondary to changes in NHERF1 RNA levels, we performed qPCR on RNA extracted from keratinocytes retrovirally transduced with vector or 16E6_WT and plated as in Fig 2A. Interestingly, NHERF1 RNA levels did not differ between keratinocytes seeded at different densities expressing either empty vector or 16E6_WT (Fig 2B).

**Fig 2.**
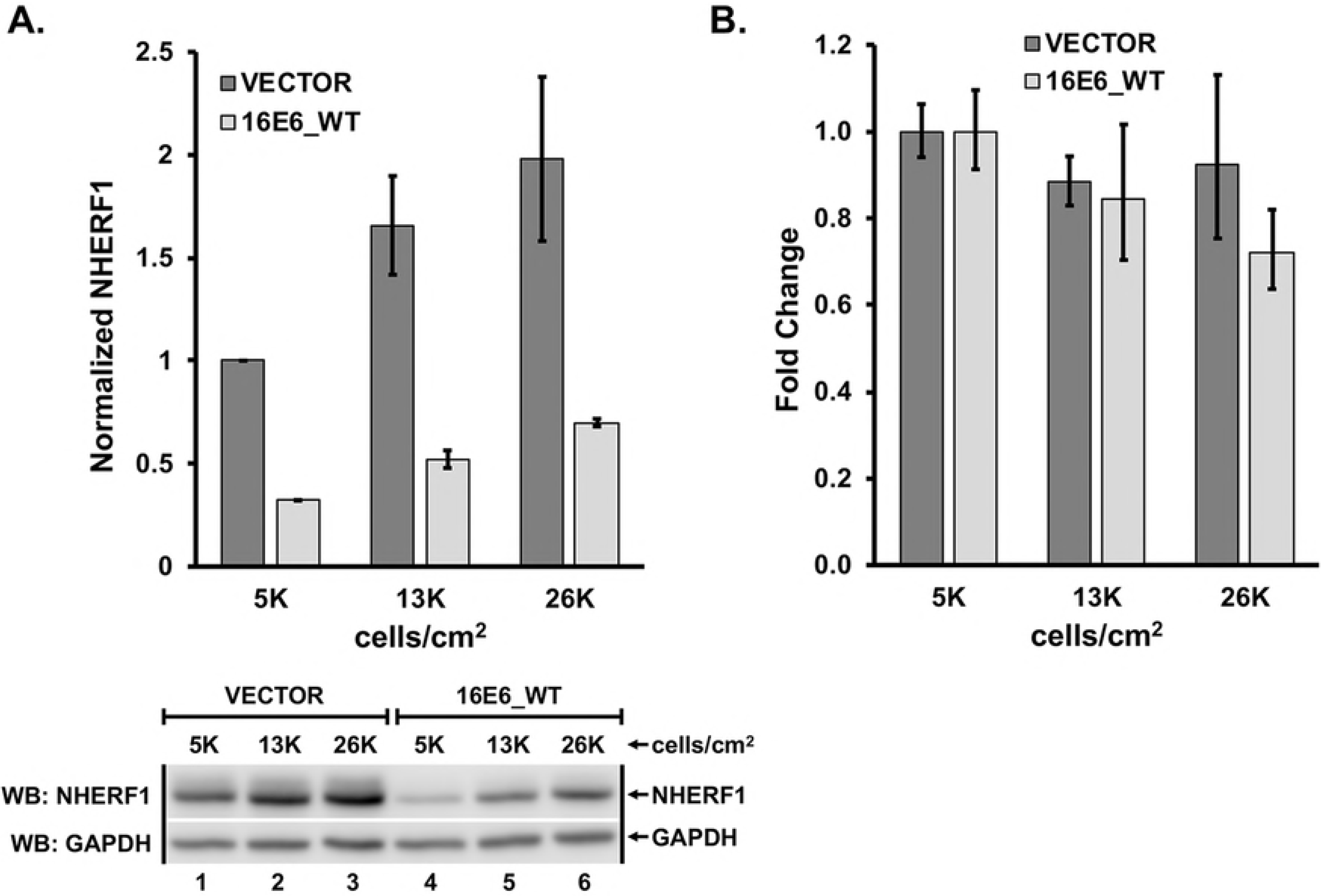
NHERF1 protein levels increase with increased cell density. (**A**) Protein levels of endogenous NHERF1 increase with cell confluency. Keratinocytes retrovirally transduced with either vector or 16E6_WT were counted and plated at the indicated cell densities. As confluency increased, NHERF1 protein levels also increased, though still reduced in the presence of 16E6_WT. The means of triplicate independent experiments ± standard error are shown. (**B**) NHERF1 RNA levels are not changed by cell confluency or by the presence of 16E6_WT. Total RNA was extracted from keratinocytes retrovirally transduced with either vector or 16E6_WT and plated at the indicated cell densities. cDNA was reverse transcribed and NHERF1 RNA levels determined by qPCR. The means of triplicate independent experiments ± standard error are shown.

### E6-mediated degradation of NHERF1 occurs via the proteasome

Because the ability of each E6 protein to reduce NHERF1 protein levels was also dependent upon the ubiquitin ligase activity of E6AP (Fig 1A), we hypothesized that E6 reduction of NHERF1 levels would be secondary to proteasome activity. We seeded retrovirally transduced keratinocytes expressing either empty vector or 16E6_WT at similar confluency and treated with either DMSO, mitomycin C (MMC) to induce p53 [30], or the proteasome inhibitor MG132 at differing concentrations for 8 hours. As expected, p53 levels increased in vector keratinocytes treated with MMC compared to untreated cells as well as in 16E6_WT cells exposed to increasing concentrations of MG132 [28] (Fig 3). NHERF1 protein levels increased significantly in a dose dependent manner upon treatment with MG132 in parallel to that seen with p53. (Fig 3, lanes 3-8). This indicated that NHERF1 is degraded through the proteasome by E6 in a manner dependent upon WT E6AP.

**Fig 3.**
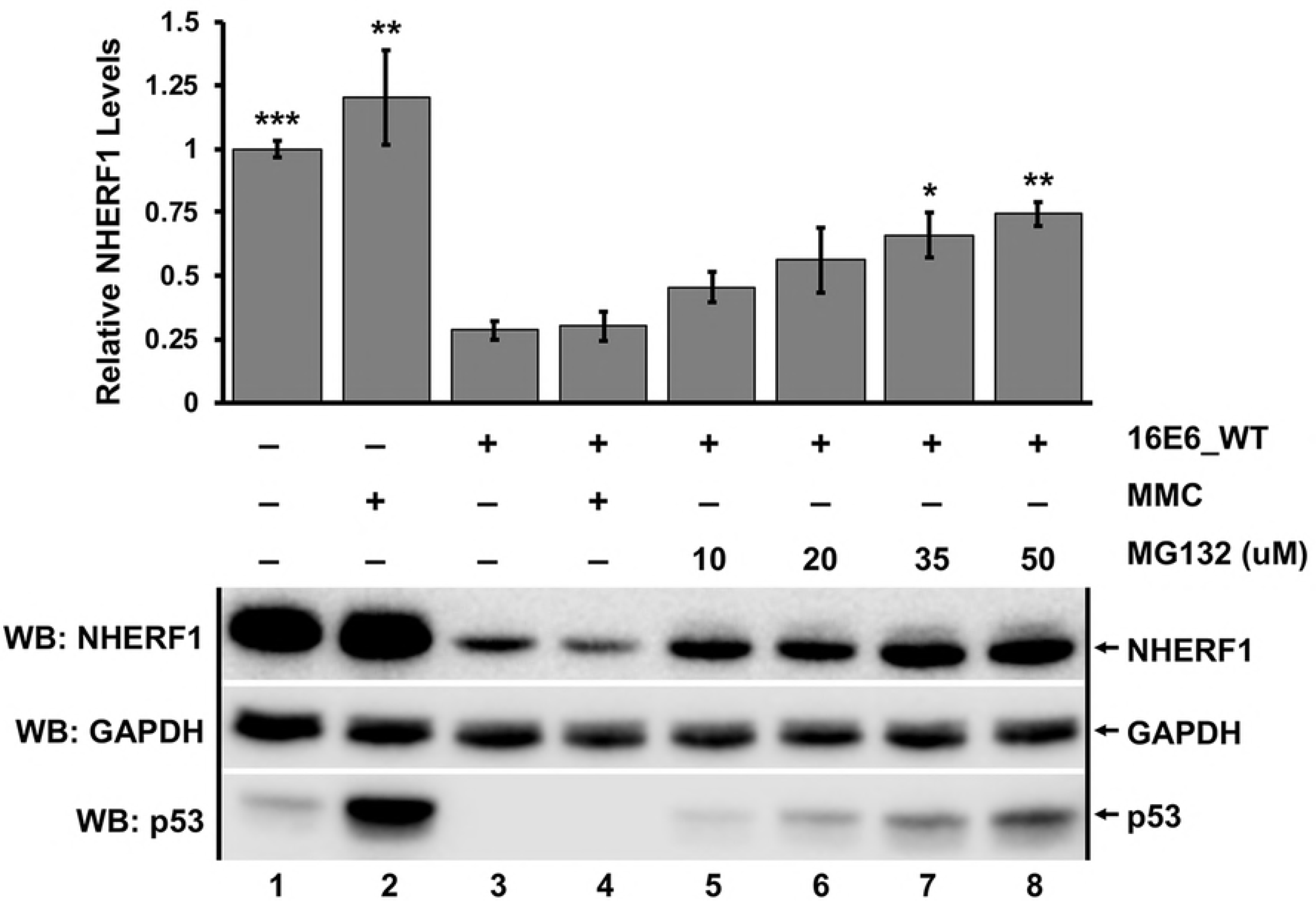
Degradation of NHERF1 by 16E6 requires proteasome function. Keratinocytes retrovirally transduced with either vector or 16E6_WT were seeded at equal confluency. Cells were treated with DMSO, mitomycin C (MMC), or the proteasome inhibitor MG132 at varying concentrations for 8 hours as indicated. MG132 significantly rescued NHERF1 protein levels in a dose dependent manner. MMC treatment was used to induce p53 levels, which were observed as a positive control. Quantification was normalized to vector-transduced cells treated with DMSO. The means of triplicate independent experiments ± standard error are shown. N=3, *<0.05, **<0.01, ***<0.001, n.s. = no significance by Student’s t-test for samples compared to untreated 16E6 keratinocytes (lane 3).

### E6-mediated degradation of NHERF1 is conserved across papillomaviruses from diverse hosts

The observation that NHERF1 was targeted by both high and low-risk HPV E6 proteins suggested that NHERF1 may also be a target of diverse non-primate E6 proteins. We examined the ability of E6 proteins from multiple different genera and different mammalian species to target NHERF1 for degradation (Fig 4). E6 proteins that preferentially bind MAML1 were unable to degrade NHERF1 (Fig 4A and 4B). All of the tested Alpha (primate), Dyodelta (boar), and Dyopi (porpoise) genera E6 proteins that bind E6AP targeted NHERF1. While E6AP-binding was necessary it was not sufficient, as E6 proteins from Omega (polar bear, UmPV1) and Omikron (cetaceans, PphPV1 and TtPV5) did not degrade NHERF1 (Fig 4A). Interestingly, E6 proteins that bind E6AP but did not target NHERF1 degradation sequence-clustered separately from E6 proteins that did target NHERF1 degradation, suggesting evolutionary divergence of this function (Fig 4B).

**Fig 4.**
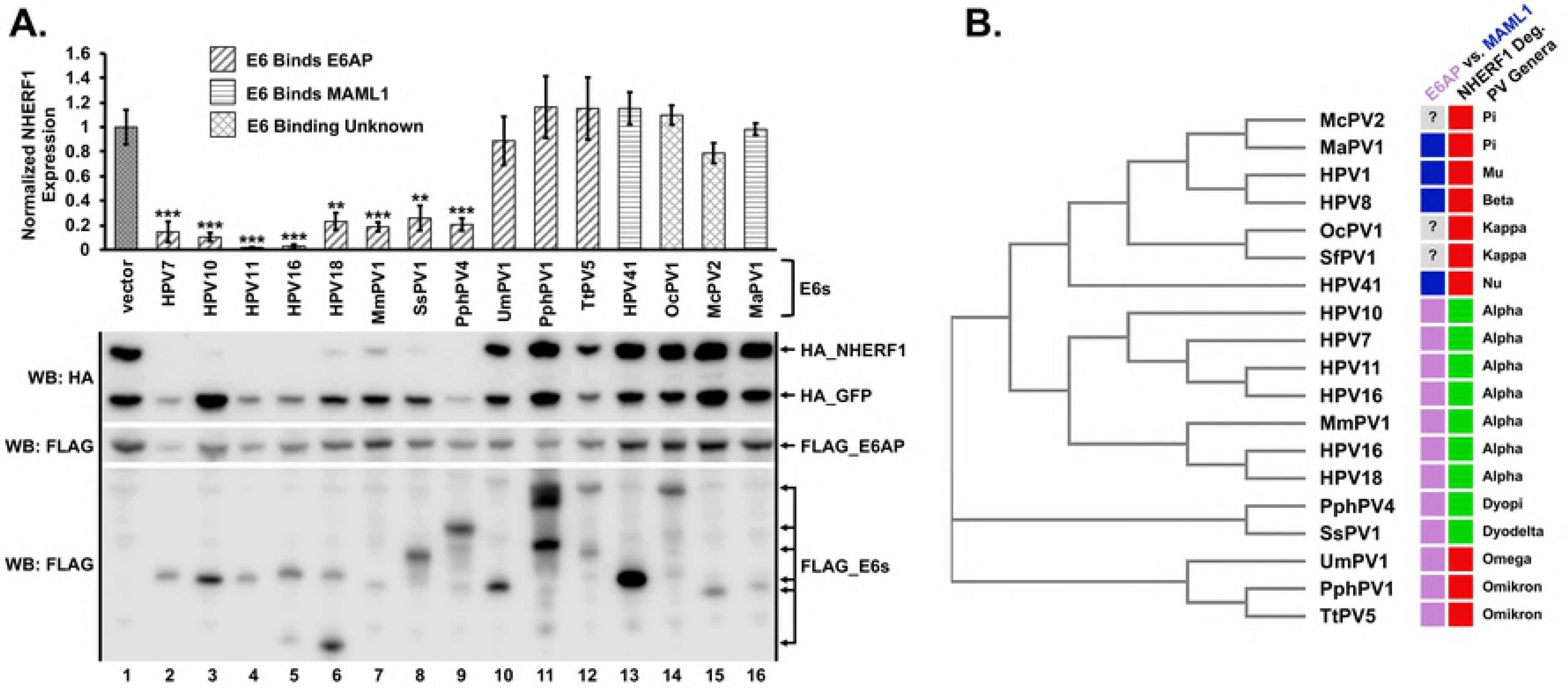
E6 proteins from evolutionarily diverse species target NHERF1. (**A**) E6 proteins from divergent animal species degrade NHERF1 via E6AP. HA_NHERF1 (0.4 ug), HA_GFP (0.1 ug), FLAG_E6AP_WT (0.35 ug), and the indicated FLAG_E6 (0.3 ug) plasmids were co-transfected into C33 cells. E6 proteins are classified based on their known preference for binding E6AP or MAML as indicated. NHERF1 was degraded by E6 proteins isolated from numerous different mammalian species. Many, but not all, of the E6 proteins that bind E6AP targeted NHERF1 for degradation, while E6 proteins that bind MAML1 did not. HA_NHERF1 protein levels in the presence of the indicated E6 proteins were normalized to co-transfected HA_GFP as an internal transfection control. A single representative blot and the means of five independent experiments ± standard error are shown. N=5, **<0.01, ***<0.001 by Student’s t-test. (**B**) E6 proteins that degrade NHERF1 cluster phylogenetically. The E6 proteins from the listed papillomaviruses were subjected to a multiple sequence alignment and then clustered phylogenetically using the program MUSCLE [65]. For E6 physical association, blue denotes MAML1 and light purple denotes E6AP. The preferential association of three E6 proteins is unknown. Ability to degrade NHERF1 is denoted in green and lack of ability to degrade NHERF1 is indicated by red. Interestingly, E6 proteins that can bind E6AP but not degrade NHERF1 cluster differently from other E6 proteins that cannot degrade NHERF1. The genera of each papillomavirus is listed. Western blot indicating NHERF1 expression in the presence of HPV1 E6, HPV8 E6, and SfPV1 E6 is shown in S3 Fig. H = *Homo sapiens* (human), Mm = *Macaca mulata* (rhesus monkey), Ss = *Sus scrofa* (wild boar), Pph = *Phocoena phocoena* (harbor porpoise), Um = *Ursus maritimus* (polar bear), Tt = *Tursiops truncatus* (bottlenose dolphin), Oc = *Oryctolagus cuniculus* (rabbit), Mc = *Mastomys coucha* (mouse), Ma = *Mesocricetus auratus* (golden hamster), Sf = *Sylvilagus floridanus* (Cottontail rabbit; CRPV1). Caption credit: Brimer N, Drews CM, Vande Pol SB. Association of papillomavirus E6 proteins with either MAML1 or E6AP clusters E6 proteins by structure, function, and evolutionary relatedness. PLoS Pathog. 2017;13(12):e1006781.

### A novel 16E6 substrate interaction domain is required for 16E6 degradation of NHERF1

Because the ability of E6 to degrade NHERF1 was not dependent upon the presence of a PBM (Figs 1 and 4), we attempted to identify which residue(s) of 16E6 were required to mediate degradation of NHERF1. The crystal structure of 16E6 complexed with the E6-binding peptide from E6AP [31] (Fig 6A) was examined to identify amino acids that were at least 20% exposed, resulting in over eighty candidate residues (S2 Fig). Candidate residues were individually mutated in the context of the 16E6 gene and the resulting point mutants were screened for their ability to degrade NHERF1 in the presence of E6AP_WT in transiently transfected C33 cells. To ensure our point mutants were not functionally defective (i.e. could not fold properly or could not interact with E6AP), we also screened the mutants for ability to degrade p53. A selection of mutants and the results of the screen are shown in Fig 5. Four mutants stood out as selectively defective in their ability to degrade NHERF1 (Fig 5B) while still being able to degrade p53 (Fig 5C): F69A, K72A, F69R and a double mutant: F69A/K72A. As evidenced in the crystal structure of 16E6, the side chains of F69 and K72 (Fig 6B) are located along the connecting alpha-helix that links the amino-terminal and carboxy-terminal zinc-structured domains of 16E6. The F69 and K72 side chains are aligned and adjacent on the connecting helix, which is on the opposite side of 16E6 from the p53 interaction surface [32] (Fig 6C).

**Fig 5.**
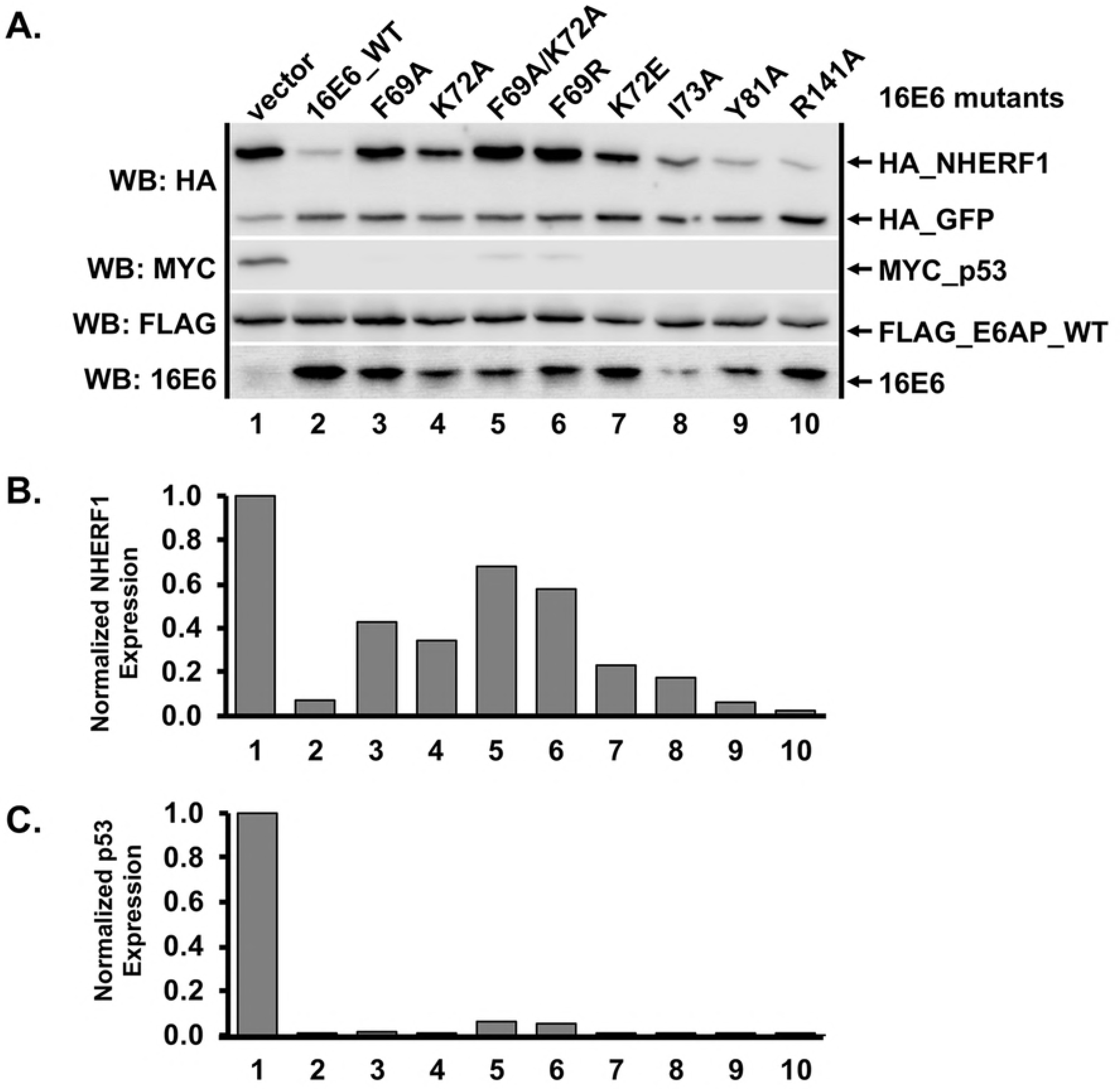
16E6 mutagenesis screen identified mutants selectively defective in their ability to degrade NHERF1. (**A**) Amino acids F69 and K72 are important for degradation of NHERF1 by 16E6. Plasmids encoding untagged 16E6_WT or 16E6 mutants (0.3 ug) were co-transfected with FLAG_E6AP (0.35 ug), HA_NHERF1 (0.4 ug), MYC_p53 (0.25 ug), and HA_GFP (0.08 ug) into C33 cells and HA_NHERF1 levels determined by western blot. Multiple 16E6 proteins were identified that were unable to degrade NHERF1 but were still capable of degrading p53. (**B**) HA_NHERF1 and (**C**) p53 protein levels were quantified and normalized to co-transfected HA_GFP as an internal transfection control.

**Fig 6.**
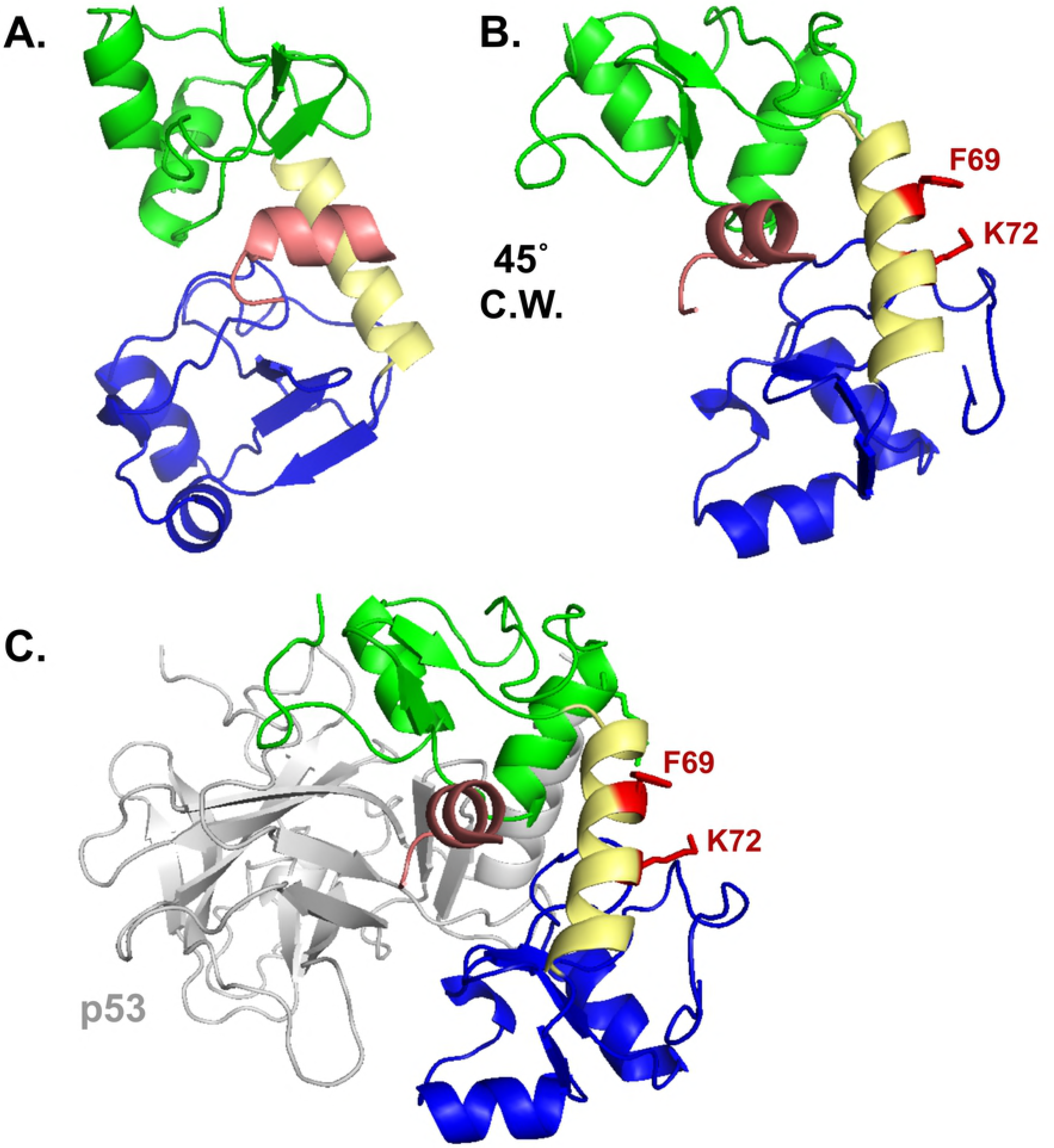
Amino acid side chains F69 and K72 define a novel substrate interaction domain on 16E6. (**A**) HPV16 E6 structure (PDB file 4GIZ) showing the amino-terminal zinc-structured domain in green, connecting alpha helix in yellow, and the carboxy-terminal zinc-structured domain in blue. The E6 protein is complexed with the LXXLL peptide of E6AP (pictured in light pink). (**B**) The E6 protein depicted in A is rotated 45° clockwise (C.W.) and the F69 and K72 residues and their side chains are highlighted in red. (**C**) A similar view as part B is shown complexed with the core p53 DNA binding domain (grey). The E6 interaction face with p53 is opposite the F69 and K72 residues.

We had identified the 16E6_F69A/K72A mutant in a transient transfection screen. To ensure the identified 16E6_F69A/K72A double mutant was selectively defective for degrading NHERF1 in the context of a stable cell line, keratinocytes retrovirally transduced with empty vector, 16E6_WT, 16E6ΔPBM, 16E6_F69A/K72A, or 11E6_WT were seeded at equal confluency and lysates prepared. Keratinocytes expressing 16E6_WT, 16E6ΔPBM, and 11E6_WT degraded NHERF1 (Fig 7, lanes 2, 3, 5). However, keratinocytes expressing 16E6_F69A/K72A were unable to stimulate the degradation of NHERF1 (Fig 7, lane 4), indicating a novel substrate interaction domain important for 16E6-mediated degradation of NHERF1.

**Fig 7.**
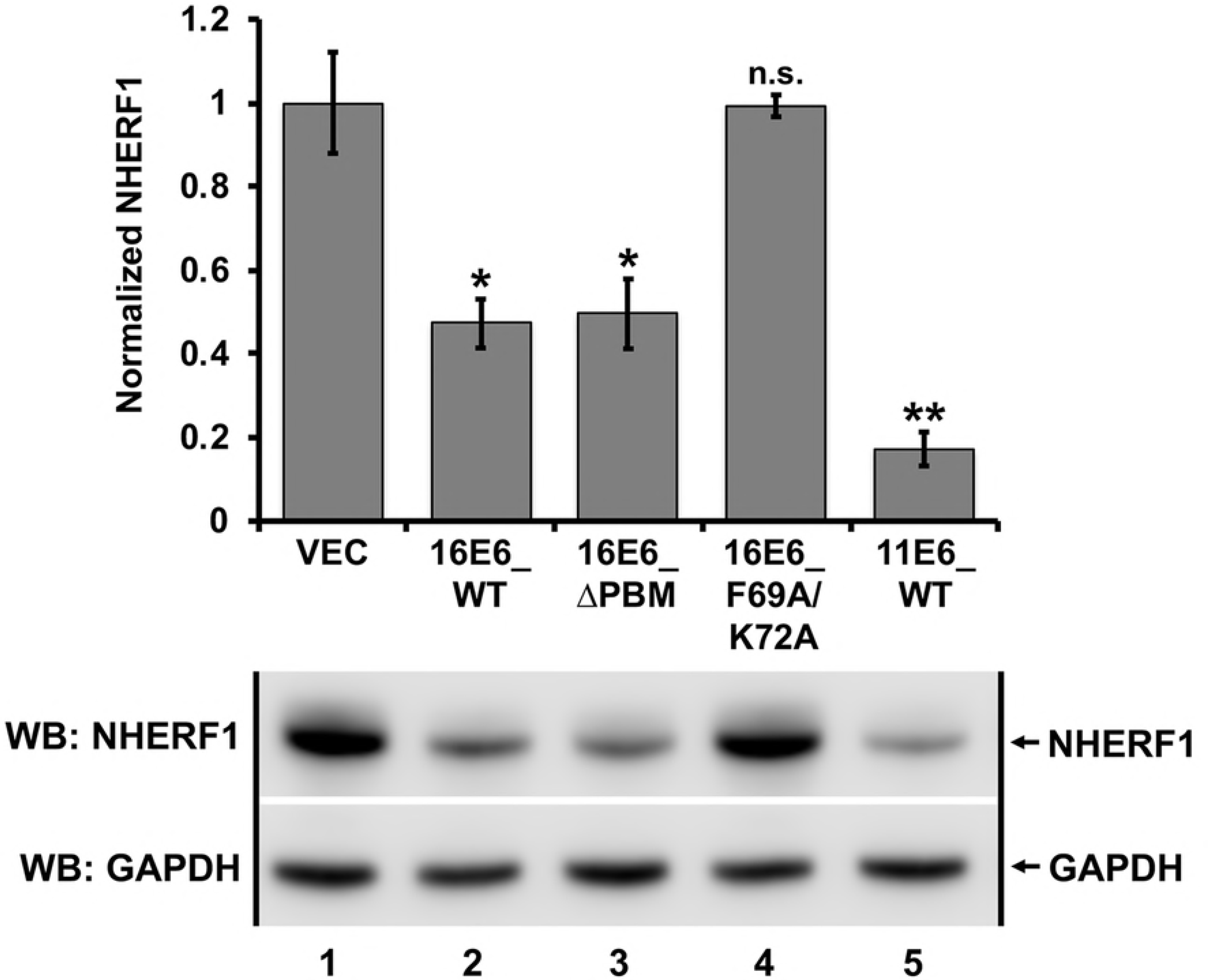
NHERF1 degradation by E6 proteins from both high and low-risk papillomaviruses in stable keratinocytes. Keratinocytes retrovirally transduced with the indicated E6 proteins were seeded at equal confluency and endogenous NHERF1 protein levels were normalized to GAPDH. 16E6_WT, 16E6 deleted of its PBM (ΔPBM), and 11E6_WT all degraded NHERF1. The 16E6_F69A/K72A double mutant did not target NHERF1 for degradation. The means of triplicate independent experiments ± standard error and one representative blot are shown. N=3, *<0.05, **<0.01, n.s. = no significance by Student’s t-test.

### Degradation of NHERF1 by 16E6 requires the NHERF1 EB domain

Because the PBM of E6 proteins is not required to initiate the degradation of NHERF1 (Figs 1, 4, and 7), we hypothesized that neither of the PDZ domains of NHERF1 would be required for 16E6 to initiate NHERF1 degradation. We truncated NHERF1 and deleted several characterized domains within the protein [33, 34] (Fig 8A). E6AP-null 8B9 cells were co-transfected with 16E6_WT, NHERF1 truncations, HA_GFP, and either E6AP_Ub^−^ or E6AP_WT. NHERF1 protein levels were quantified, and then normalized to the internal transfection control (HA_GFP). The various NHERF1 truncations displayed different expression levels. To account for these variations, levels of NHERF1 truncations in the presence of E6AP_Ub^−^ were set to 100% and the expression level of the corresponding NHERF1 truncation in the presence of E6AP_WT was normalized accordingly (Fig 8B and 8C, bar graphs). All NHERF1 truncations containing the EB domain were targeted for degradation by 16E6 in the presence of E6AP_WT (highlighted in green in Fig 8A). Truncations of NHERF1 that lacked the EB domain were not targeted for degradation by 16E6 (highlighted in red in Fig 8A). In addition, the NHERF1 PBM was not required for 16E6 mediated degradation (Fig 8C, lanes 5 vs. 6 and 9 vs. 10).

**Fig 8.**
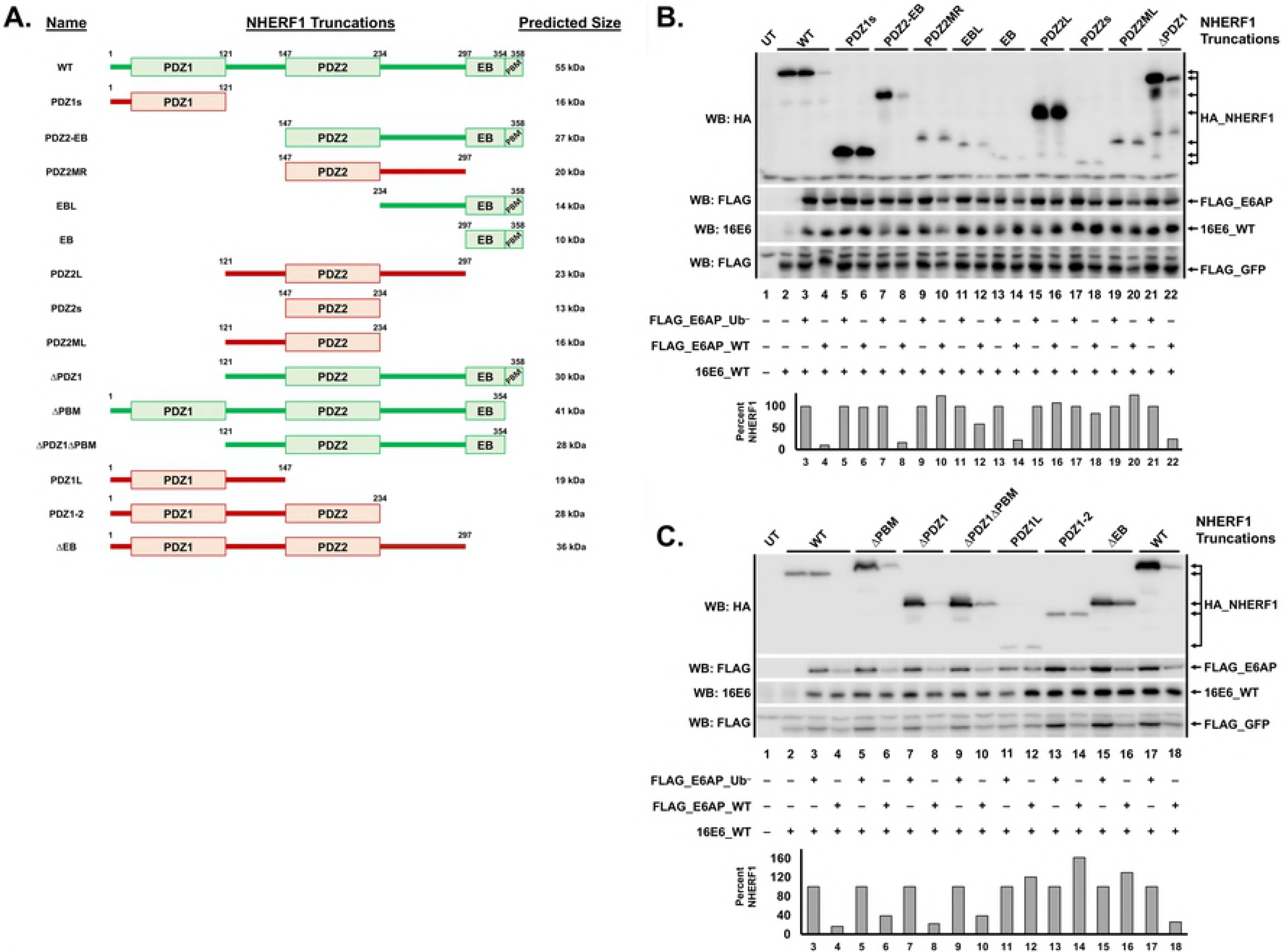
NHERF1 truncations identify the EB domain as necessary for NHERF1 degradation by 16E6. (**A**) Schematic of NHERF1 truncations. NHERF1 proteins that were successfully degraded by 16E6_WT are depicted in green while truncations that were not degraded are depicted in red. (**B** and **C**) NHERF1 truncations containing the EB domain were degraded, while those lacking the EB domain were not. The listed HA_NHERF1 truncations (shown in A in the order loaded in B and C, 0.8 ug), untagged 16E6_WT (1 ug), FLAG_GFP (0.08 ug), and either FLAG_E6AP_WT (1.2 ug) or FLAG_E6AP_Ub^−^ (1.2 ug, defective for ubiquitin ligase activity) were co-transfected in E6AP-null 8B9 cells. HA_NHERF1 levels were quantified and normalized to FLAG_GFP as an internal transfection control. The bar graph below the blot represents quantification of each listed HA_NHERF1 truncation. In panel C, the WT NHERF1 in lanes 2-4 contains an amino terminal 1X HA tag while the WT NHERF1 in lanes 17 and 18 contains an amino terminal 2X HA tag. All of the NHERF1 truncations contain amino terminal 2X HA tags. Levels of HA_NHERF1 truncations in the presence of FLAG_E6AP_WT were normalized to their corresponding expression in the presence of FLAG_E6AP_Ub^−^ to account for the differing expression levels. UT = untransfected.

We identified the NHERF1 EB domain as a requirement for 16E6 mediated degradation and the importance of 16E6 residues F69 and K72. In order to examine the interactions between the 16E6+E6AP+NHERF1 complex, all three proteins were expressed in a yeast three-hybrid system so as to detect the heterotrimeric complex. We fused 16E6_WT and ubiquitin ligase dead E6AP (E6AP_Ub^−^) to the LexA DNA binding domain and co-expressed this fusion with either vector, 16E6_WT, or 16E6_F69A/K72A in yeast containing a LexA responsive LacZ reporter. These yeast were then mated to yeast expressing native p53 or Gal4 (G4) transactivator fusions to NHERF1 121-358 (containing the EB domain), NHERF1 121-297 (deleted of the EB domain), 16E6_WT, or the tyrosine phosphatase PTPN3 (a PDZ protein) (Fig 9). The LexA_16E6 fusion co-expressed with p53 (in the absence of E6AP) resulted in very weak activation of the LacZ reporter (spot 4B) while co-expression with G4_PTPN3 resulted in strong transactivation (spot 6B), but no interaction with NHERF1 (spots 2B and 3B). We then co-expressed 16E6 and E6AP by using a LexA_E6AP_Ub^−^ fusion together with native 16E6. When LexA_E6AP_Ub^−^, untagged 16E6_WT, and p53 were co-expressed, a strong activation of the LacZ reporter was observed (Fig 9, spot 4D), illustrating that while p53 has a weak direct interaction with 16E6, it interacts strongly with 16E6 bound to E6AP. This activation was also seen with 16E6_F69A/K72A in the presence of LexA_E6AP_Ub^−^ and p53 (Fig 9, spot 4E), indicating the preserved ability of the 16E6_F69A/K72A double mutant to bind E6AP and recruit p53. When LexA_E6AP_Ub^−^, 16E6_WT, and G4_NHERF1 121-358 (contains the EB domain) were co-expressed, we observed activation of the LacZ reporter, indicating the recruitment of NHERF1 to E6AP by 16E6_WT (Fig 9, spot 2D). Truncating the EB domain from the G4_NHERF1 (G4_NHERF1 121-297, spot 3D) or the use of the 16E6_F69A/K72A double mutant (spot 2E) ablated the reporter transactivation, indicating the requirement of the EB domain and the importance of 16E6 residues F69 and K72 in the interaction of the 16E6+E6AP+NHERF1 complex.

**Fig 9.**
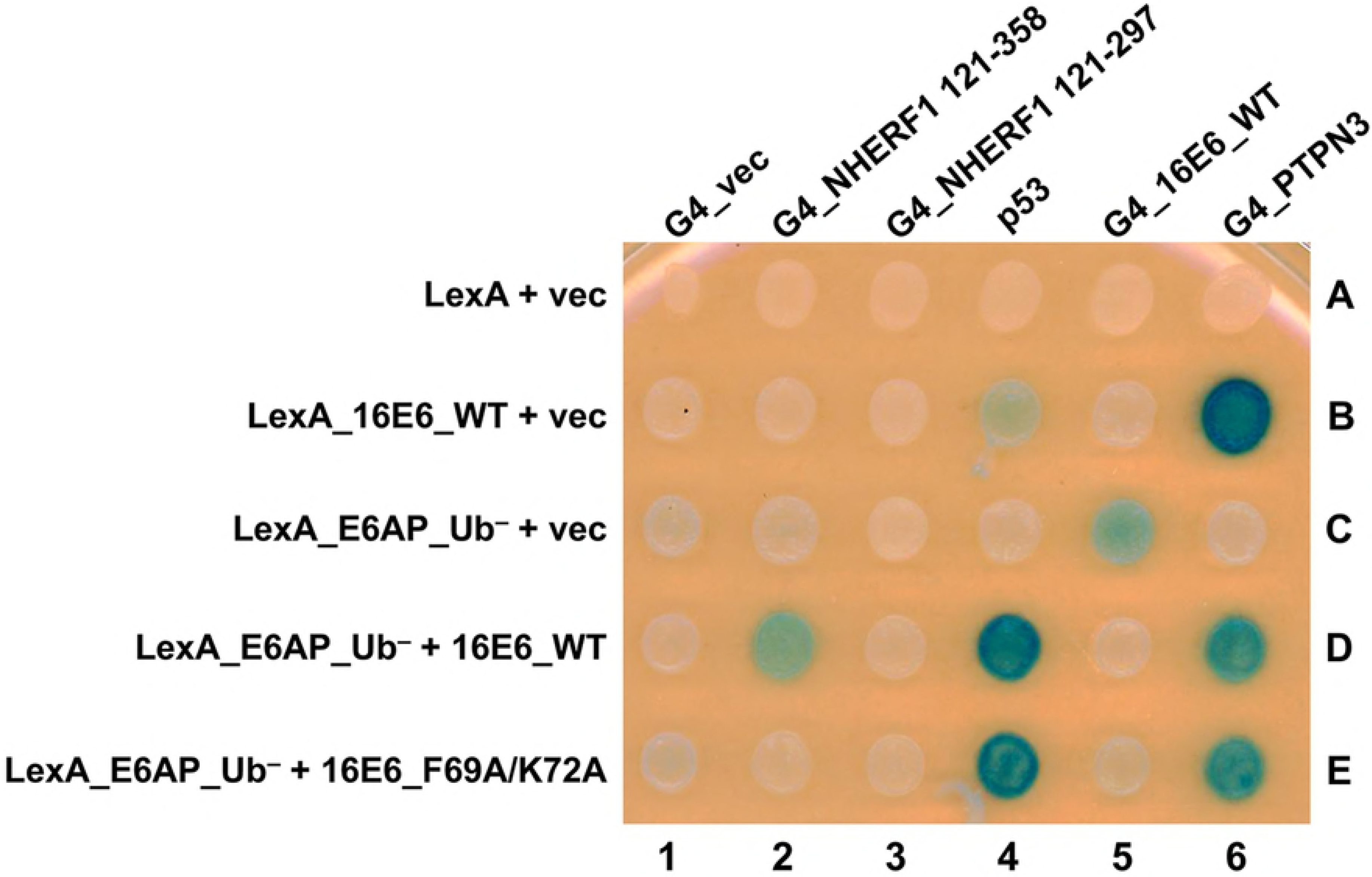
The E6-E6AP-NHERF1 complex can be modeled in yeast. Yeast three-hybrid plasmids expressing the LexA DNA binding domain fused to either 16E6_WT or E6AP_Ub^−^ were co-expressed in yeast (bait) together with either vector, 16E6_WT, or 16E6_F69A/K72A as indicated. The bait yeast were mated to prey yeast expressing Gal4 activation domain (G4), or G4 fused to 16E6_WT, PTPN3, truncations of NHERF1, or native p53 and diploids selected. Positive controls for 16E6 expression included the established interaction of the 16E6 PBM with the PDZ domain of tyrosine phosphatase PTPN3 and 16E6-E6AP complex interaction with p53. 16E6_WT recruited NHERF1, p53, and PTPN3 to LexA_E6AP_Ub^−^. The recruitment of NHERF1 to LexA_E6AP_Ub^−^ by 16E6 was specifically lost upon mutation of residues F69 and K72, however, p53 and PTPN3 recruitment were maintained. 16E6_WT recruitment of NHERF1 was not seen with an NHERF1 truncation lacking the EB domain (G4_NHERF1 121-297). Caption credit: Ansari T, Brimer N, Vande Pol SB. Peptide interactions stabilize and restructure human papillomavirus type 16 E6 to interact with p53. J Virol. 2012;86(20):11386-91.

### E6AP-dependent NHERF1 degradation by E6 activates the canonical Wnt/β-catenin pathway

It has been shown that high-risk HPV E6 proteins augment the canonical Wnt/β-catenin signaling pathway [35–39]. Additionally, it has been shown that NHERF1 inhibits the canonical Wnt/β-catenin signaling pathway through multiple mechanisms. NHERF1 forms a complex with β-catenin [40] and can also bind to the intracellular PBM of certain isoforms of Frizzled [41], a G-protein coupled receptor important in the activation of the canonical Wnt signaling pathway. Therefore, we hypothesized E6 degradation of NHERF1 would activate the Wnt/β-catenin signaling pathway in cells expressing E6. To test this possibility, we utilized the TOP/FOP luciferase reporter assay. 16E6, 16E6ΔPBM, 11E6, and 18E6 all stimulated the activity of the Wnt/β-catenin pathway over vector-transfected cells (Fig 10). However, cells transfected with 16E6_F69A/K72A were unable to augment the canonical Wnt pathway over vector levels, indicating that the ability of E6 to degrade NHERF1 is required for E6 activation of the canonical Wnt/β-catenin signaling pathway.

**Fig 10.**
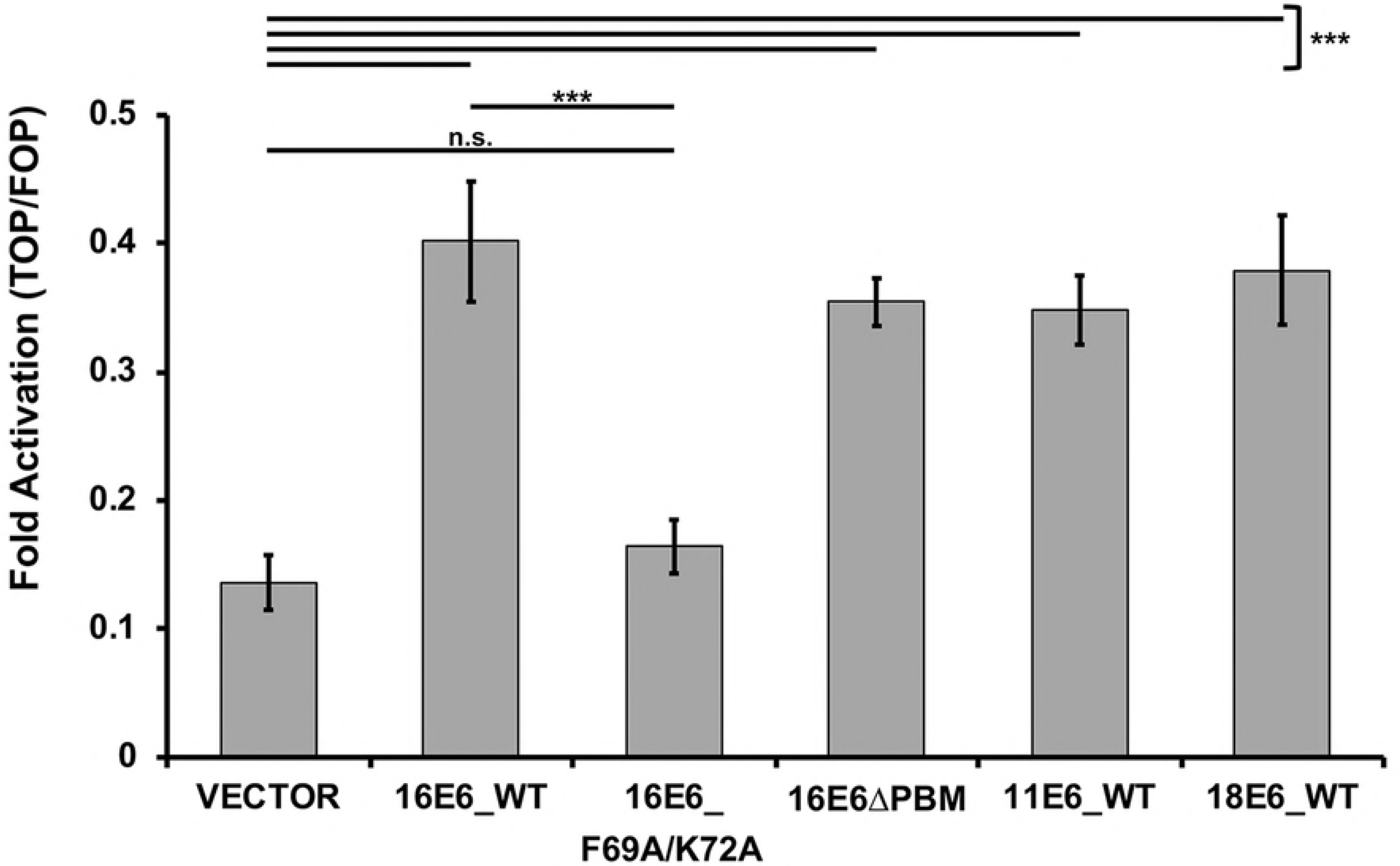
Activation of the canonical Wnt/β-catenin pathway is augmented by E6 proteins that can degrade NHERF1. The listed E6 proteins were co-expressed with FLAG_E6AP_WT, the TOPFLASH or FOPFLASH luciferase reporter, and a renilla luciferase internal transfection control plasmid in C33A cells. Transfected cells were treated with Wnt3A conditioned media for 8.5 hours, lysed in 1X passive lysis buffer (Promega), and measured for luciferase and renilla luminescence. Fold activation was determined by normalizing the TOPFLASH luminescence by the FOPFLASH luminescence. Each E6 protein that could degrade NHERF1 (16E6_WT, 16E6_ΔPBM, 11E6_WT, and 18E6_WT) augmented the canonical Wnt pathway. 16E6_F69A/K72A, which cannot degrade NHERF1, failed to increase Wnt pathway activation over vector levels. Statistical significance was determined from three independent experiments by Student’s t-test (***<0.001, n.s. = no significance).

The earlier Accardi et al. study proposed that expression of 16E7 sensitized NHERF1 for degradation by the induction of NHERF1 phosphorylation [26]. Our experiments did not show either E7 induction of slow-migrating NHERF1 phosphorylated isoforms or an enhancement of E6-NHERF1 degradation upon co-expression of E7 (S3 and S4 Figs).

## Discussion

E6 proteins from papillomaviruses can be separated into distinct groups: those that bind MAML1 and repress Notch signaling, and those that bind E6AP and hijack its ubiquitin ligase activity [7, 42–44]. E6 proteins from papillomaviruses in the Alpha, DyoDelta, Dyopi, Omega, and Omikron genera all behave similarly in that they bind E6AP and activate its ubiquitin ligase activity [7]. Here, we describe the degradation of NHERF1 by E6 proteins from both high and low-risk HPVs, as well as from papillomaviruses from multiple divergent mammalian species. The ability of these E6s to degrade NHERF1 is dependent upon E6AP (Fig 1) and the proteasome (Fig 3). In addition, we identify two amino acids in 16E6 (F69 and K72) that are necessary for E6-mediated degradation of NHERF1. These two residues are aligned, and adjacent in the outwardly oriented face of the E6 connecting alpha helix, suggesting a novel interaction domain (Fig 6B and 6C). NHERF1 degradation by E6 requires the NHERF1 EB domain, but does not require the PBM at the extreme carboxy terminus of NHERF1 (Fig 8B and 8C). The ability of E6 proteins to degrade NHERF1 augments the canonical Wnt/β-catenin signaling (Fig 10), an oncogenic pathway frequently active in cancer.

NHERF1 is the product of the SLC9A3R1 gene. SLC9A3R1 mRNA is broadly expressed in epithelia, with the highest mRNA expression in kidney, gut, and esophagus. NHERF1 is not developmentally essential, although mice have considerably reduced lifespans [45]. NHERF1-null mice are prone to phosphate wasting, brittle bone structure, and hydrocephaly [45] due to the mislocalization of proteins with which NHERF1 normally associates [45–47]. NHERF1 contains two PDZ domains and an EB domain at the carboxy terminus through which it interacts with ezrin, radixin, and moesin to link itself, and proteins to which it is bound, to the actin cytoskeleton network [48]. While the functions of NHERF1 are varied due to its role as a scaffold, multiple studies indicate it regulates cell growth and differentiation, two key cellular functions that papillomaviruses disrupt in the process of viral infection.

Whether NHERF1 is a tumor suppressor or an oncogene has been debated in the literature. There are numerous papers regarding NHERF1 human cancer phenotypes, but they are collectively inconsistent [49, 50]. NHERF1-null mice do not have a direct cancer phenotype, but have lengthened intestines [51], indicating a growth regulatory function of NHERF1. The diminished life span of NHERF1 null mice could limit observation of cancer traits. However, a recent in vivo study provided strong genetic support for NHERF1 as a tumor suppressor. APC^Min/+^ mice bred as either heterozygote or knockout for NHERF1 experience considerably shorter survival than their NHERF1-expressing counterparts due to increased tumor burden, demonstrating a tumor suppressor phenotype for NHERF1 [51]. Additionally, these APC^Min/+^ mice lacking NHERF1 have greater activation of Wnt/β-catenin signaling, suggesting NHERF1 acts as a negative regulator of this oncogenic pathway. NHERF1-associated proteins that plausibly could regulate cell proliferation are numerous and include β-catenin [40], Frizzled [41], G-protein coupled receptors (β-adrenergic type 2, [52]), receptor tyrosine kinases (PDGFR, [53]), phosphatases (PTEN, [54]), transcriptional coactivators (YAP1, [55]), ion channels (Kir1.1 and CFTR, [56]), phospholipase-C [57], and actin anchoring proteins (ezrin, radixin, and moesin, [48]).

Several studies have indicated that HPV E6 proteins can activate canonical Wnt/β-catenin signaling [35–39]. Our work expands and builds upon the scope of these studies. The ability of E6 to degrade NHERF1 and activate Wnt signaling may aid in propagation of papillomaviruses by enhancing the stimulation of cellular proliferation and promoting cell survival. There are numerous cell growth regulatory avenues that E6 could manipulate by degrading NHERF1 and within this study we have explored one possibility: the canonical Wnt/β-catenin signaling pathway (Fig 10); other possibilities will be the subject of future studies.

The EB domain of NHERF1 is required for E6-mediated degradation in the presence of E6AP (Fig 8B and 8C). This domain is responsible for linking NHERF1 to the actin cytoskeleton network via interaction with ERM proteins [48]. NHERF1 has a PBM at its extreme carboxy terminus and when the EB domain is not bound to ERM proteins, the NHERF1 PBM can self-associate with the NHERF1 PDZ2 domain, resulting in a closed NHERF1 conformation [33]. The head-to-tail closed NHERF1 confirmation is not required for E6-mediated degradation, as an NHERF1ΔPBM mutant was still targeted for degradation by E6 in the presence of E6AP_WT (Fig 8C). Nor was the 16E6 PBM required for degradation of NHERF1 (Figs 1, 4, and 7), contrary to a prior report [26].

In addition to the requirement of the NHERF1 EB domain, two E6 residues, F69 and K72, are necessary for E6-mediated NHERF1 degradation (Figs 5 and 7). Crucially, the 16E6_F69A/K72A double mutant can still initiate the degradation of p53, indicating it is still able to bind E6AP, recruit p53 to the complex, and trigger ubiquitination. The F69 and K72 residues are also required to form a tri-molecular complex between E6AP, E6, and NHERF1 in yeast (Fig 9, spot 2E vs. 2D). Like the association of E6 with p53, NHERF1 does not interact directly with E6, but requires prior association of E6 with E6AP, indicating that NHERF1 requires an altered conformation of E6 that is secondary to E6 binding to E6AP [58].

As we were testing the ability of E6 proteins to degrade NHERF1 in stable keratinocyte cell lines, we discovered that NHERF1 protein levels are sensitive to cell confluency (Fig 2A). The relationship between NHERF1 and cell confluency may contribute to the lack of identification of NHERF1 as a degradation target of low-risk E6 proteins in the past, as well as differences between our studies and a prior publication [26]. It is likely that this observation underlies disparate findings between different laboratories regarding NHERF1 cancer associated traits [49, 50]. Future studies of NHERF1 must take into account and carefully control cell densities when performing experiments.

Binding to E6AP is necessary for E6-induced degradation of NHERF1, but it is not sufficient, as three tested E6 proteins that bind E6AP do not target NHERF1 for degradation: UmPV1 E6 (polar bear), PphPV1 E6 (porpoise), and TtPV5 E6 (bottlenose dolphin) (Fig 4A). Interestingly, the three E6 proteins that do not degrade NHERF1 cluster together in phylogenetic relatedness (Fig 4B). We utilized transfected human NHERF1 throughout our study, so it is possible that the inability of these three E6 proteins to target NHERF1 for degradation may be due to evolutionary divergence in the NHERF1 homologs. Future studies will explore if the lack of degradation of human NHERF1 by UmPV1, PphPV1, and TtPV5 is due to evolutionary divergence of the respective NHERF1 proteins compared to human NHERF1. It would be of interest to determine if NHERF1 is a “universal” target of E6 proteins that act through association with E6AP.

Discovery of NHERF1 as a novel target for not only high and low-risk mucosal and cutaneous HPV E6 as well as a wide range of E6 proteins across divergent host species indicates a significant and previously undescribed role for NHERF1 in papillomavirus biology. That NHERF1 is a conserved target of papillomavirus E6 proteins further elevates the importance of NHERF1 as a cell growth regulator. Finally, the identification of this highly conserved E6 degradation target may represent a novel avenue for therapeutic intervention against both low and high-risk HPV.

## Materials and methods

### Cells and cell culture

E6AP-null 8B9 cells (a gift of Dr. Lawrence banks, ICGEB, Italy) [59] and HPV-negative C33A cervical cancer cells (ATCC) were maintained and transfected using polyethylenimine (PEI) as previously described [58]. Normal immortalized keratinocytes (NIKS, obtained from ATCC) are spontaneously immortalized foreskin keratinocytes [60] that were cocultured with mitomycin C-treated 3T3 feeder cells in F medium as described previously [61]. NIKS were retrovirally transduced with replication-defective murine retroviruses based on pLXSN [62] as previously described [25]. Retrovirally transduced NIKS cells were counted and seeded at equal confluency in each experiment.

### Plasmids

Epitope tagged E6AP, GFP, E6, and NHERF1 were all transiently expressed from the pcDNA3 plasmid. HA-tagged NHERF1 originated from Vijaya Ramesh’s laboratory (from Addgene, plasmid 11635). 16E6 point mutants were created using QuikChange primer design (Agilent Technologies). NHERF1 truncations were PCR generated and sequenced.

### Antibodies and Western blots

12 well plates of transfected mammalian cells were lysed in 0.5X IPEGAL as described previously [7]. Transduced NIKS were lysed in 1% SDS, 5mM EDTA, and 1 mM sodium vanadate and equilibrated for protein content (Biorad assay kit). All lysates were resolved by SDS-PAGE electrophoresis and transferred to PVDF membranes. Antibodies: anti-HA (Bethyl Laboratories, Inc.), anti-FLAG M2 (Sigma), anti-p53 Ab-8 (ThermoFisher Scientific), anti-16E6 6G6 (a generous gift from Arbor Vita Corporation), anti-SLC9A3R1 (Sigma), anti-GAPDH (Cell Signaling Technology), and anti-MYC 9B11 (Cell Signaling Technology).

### RT-PCR

Retrovirally transduced NIKS were plated at different cell densities and harvested following a TRIzol RNA harvest protocol (Invitrogen). cDNA was generated using random hexamers. Quantitative real-time PCR was performed on the cDNA using iQ™ SYBR^®^ Green Supermix (BioRad #1708880). The primers targeted the SLC9A3R1 gene (BioRad Assay ID: qHsaCEP0050521) and the GAPDH gene (BioRad Assay ID: qHsaCEP0041396). Relative values were analyzed using the ΔΔC_⊤_ method (where C_⊤_ is the threshold cycle) and GAPDH as a control.

### Wnt/β-catenin luciferase reporter assay

C33A cells plated at 70% confluency were transiently transfected with DNA of the TOPFLASH or control FOPFLASH (containing mutated TCF/β-catenin binding sites; 1 ug) plasmid, Renilla luciferase (0.005 ug) plasmid (used to evaluate transfection efficiency), FLAG_E6AP_WT (0.35 ug) plasmid, and the indicated E6 plasmids (0.3 ug). 18 hrs post-transfection, media was removed and Wnt3A conditioned media was added for 8.5 hours to stimulate the Wnt pathway. Luciferase levels were measured using the Dual-Luciferase^®^ Reporter Assay System (Promega) and a Cytation1 Plate Reader (software version 3.04.17). FOPFLASH luciferase readings were low, and were subtracted from the paired TOPFLASH readouts. 10% fetal bovine serum Wnt3A conditioned media was generated using L Wnt-3A murine fibroblasts (ATCC, CRL-2647) as previously described [63].

### Phylogenetic analysis

Multiple protein sequence files were downloaded from the Papillomavirus Episteme [64] and aligned using the EMBL-EBI MUSCLE (<u>MU</u>ltiple <u>S</u>equence <u>C</u>omparison by <u>L</u>og-<u>E</u>xpectation) program [65]. The phylogenetic tree was generated as a neighbour-joining tree without distance corrections within the MUSCLE program [65].

### Yeast expression

Modified LexA-based yeast three-hybrid assays were performed as previously described [58].

## Acknowledgements

We thank Nicole Brimer for assistance with technical challenges that were encountered throughout this study and Kelly Drews for extensive discussions and for critical reading of the manuscript.

## Supporting information legends

**S1 Fig. Reduction of NHERF1 protein levels is not an overexpression artifact.** Titrations of the indicated three different E6 proteins (16E6_WT, 16E6_ΔPBM, and 11E6_WT) were co-transfected with FLAG_E6AP_WT (1 ug), HA_GFP (0.02 ug), and either HA_NHERF1 (0.5 ug) or p53 (0.5 ug) in E6AP-null 8B9 cells. A representative blot of the triplicate experiments for each E6 protein is shown. Increased E6 expression for 16E6_WT, 16E6ΔPBM, and 11E6_WT resulted in decreased NHERF1 protein levels. Both 16E6_WT and 16E6ΔPBM degrade p53 with increasing E6 expression. Overexpression of 11E6 _WT (>0.1 ug E6) resulted in degradation of co-expressed E6AP_WT.

**S2 Fig. 16E6 point mutants screened to determine amino acid(s) necessary for NHERF1 degradation.** The 16E6 crystal structure (PDB file 4GIZ) was examined for residues that were at least 20% exposed as determined by the Swiss PDB Viewer. Point mutants of these identified amino acids were then screened to identify which residue(s) resulted in an E6 protein that was selectively defective for degrading NHERF1 but retained degradation of p53. Residues of interest are indicated in red.

**S3 Fig. The E7 papillomavirus protein does not induce phospho-specific isoforms of NHERF1.** Keratinocytes retrovirally transduced with vector or the indicated E7 and/or E6 proteins were seeded at equal confluency. Levels of endogenous NHERF1 were determined by western blot. Levels of phosphorylated NHERF1 (pNHERF1) were unchanged in keratinocytes expressing empty vector compared to the various E7 proteins. Keratinocytes expressing the E6 protein from high-risk (HPV16) and low-risk (HPV11) degraded NHERF1. H = *Homo sapiens* (human), Sf = *Sylvilagus floridanus* (Cottontail rabbit; CRPV1). Caption credit: Accardi R, Rubino R, Scalise M, Gheit T, Shahzad N, Thomas M, et al. E6 and E7 from human papillomavirus type 16 cooperate to target the PDZ protein Na/H exchange regulatory factor 1. J Virol. 2011;85(16):8208-16.

**S4 Fig. The presence of the E7 oncoprotein does not enhance NHERF1 degradation by E6 proteins.** C33A cells were co-transfected with the following plasmids: HA_NHERF1 (0.4 ug), FLAG_E6AP_WT (0.35 ug), HA_GFP (0.08 ug), the indicated E6 protein (0.3 ug), and the indicated E7 protein (0.3 ug). HA_NHERF1 levels were determined by western blot. FLAG_18E6* is a truncated splice isoform of 18E6. Quantitation is derived from three experimental replicates. A representative blot and means of triplicate independent experiments ± standard error are shown. N=3. **<0.01, ***<0.001, ****0.0001, n.s. = no significance by Student’s t-test.

